# *De Novo* TANGLED1 Recruitment to Aberrant Cell Plate Fusion Sites in Maize

**DOI:** 10.1101/2024.03.07.583939

**Authors:** Aimee N. Uyehara, Beatrice N. Diep, Lindy A. Allsman, Sarah G. Gayer, Stephanie E. Martinez, Janice J. Kim, Shreya Agarwal, Carolyn G. Rasmussen

## Abstract

Division plane positioning is critical for proper growth and development in many organisms. In plants, the division plane is established before mitosis, by accumulation of a cytoskeletal structure called the preprophase band (PPB). The PPB is thought to be essential for recruitment of division site localized proteins, which remain at the division site after the PPB disassembles. Here, we show that a division site localized protein, TANGLED1 (TAN1), is recruited independently of the PPB to the cell cortex at sites, by the plant cytokinetic machinery, the phragmoplast. TAN1 recruitment to de novo sites on the cortex is partially dependent on intact actin filaments and the myosin XI motor protein OPAQUE1 (O1). These data imply a yet unknown role for TAN1 and possibly other division site localized proteins during the last stages of cell division when the phragmoplast touches the cell cortex to complete cytokinesis.

**Summary Statement:** The plant division site protein TANGLED1 is recruited to *de novo* cell plate insertion sites independently of the preprophase band.

## Introduction

Asymmetric or formative cell divisions generate daughter cells with different cell fates. In Arabidopsis, stomata are generated through a series of asymmetric divisions initiated by basic helix-loop-helix transcription factor mediated signaling and reinforced by polarized protein accumulation (Bergmann and Sack, 2007; Dong and Bergmann, 2010; Pillitteri and Torii, 2012; Putarjunan *et al*., 2019; Rowe *et al*., 2019; Gong *et al*., 2021; Seo *et al*., 2022; Smit and Bergmann, 2023). Many of the same signaling networks are required for grass stomatal development, although some proteins are repurposed for specialized divisions (Raissig *et al*., 2017; Wu *et al*., 2019; Hughes and Langdale, 2022). The grass-specific stomatal complex contains additional cells that flank the guard cells called subsidiary cells. An invariantly localized asymmetric division generates the small, crescent shaped subsidiary cell and a larger pavement cell (Facette *et al*., 2015; Gray, Liu and Facette, 2020; Durney *et al*., 2023). Subsidiary cell division positioning is an excellent system due to the consistently positioned divisions and its well-characterized signaling pathways (Frank, Cartwright and Smith, 2003; Cartwright, Humphries and Smith, 2009; Humphries *et al*., 2011; Zhang *et al*., 2012, 2022; Sutimantanapi, Pater and Smith, 2014; Raissig *et al*., 2017; H. Wang *et al*., 2019; Wu *et al*., 2019; Li *et al*., 2021; Ashraf, Liu and Facette, 2023; Durney *et al*., 2023; Nan, Char, *et al*., 2023; Nan, Liang, *et al*., 2023).

Two cytoskeletal structures participate in division plane positioning: the PPB, which assembles during late G2, and the phragmoplast, which assembles during telophase and expands to complete cytokinesis. The PPB is a cortical ring of microtubules and actin that is an early indicator of the location of cell division that disassembles before metaphase (Pickett-Heaps and Northcote, 1966; Kakimoto and Shibaoka, 1987; Mineyuki, 1999; Smertenko *et al*., 2017). Following chromosome and organelle redistribution in metaphase and anaphase, the phragmoplast forms to facilitate cell plate expansion. The cell plate fuses with the mother cell plasma membrane to divide the two daughter cells (Gunning, 1982; Samuels, Giddings and Staehelin, 1995; Müller and Jürgens, 2016). If the cell plate fuses at the location previously marked by the PPB, the location is called the division site (Smertenko *et al*., 2017). The more general location where the cell plate fuses, whether at the division site or in another location, is called the cell plate fusion site (Smertenko *et al*., 2017).

Genetic disruption of PPB formation often leads to significantly stunted growth and division plane positioning defects (Torres-Ruiz and Jürgens, 1994; Camilleri *et al*., 2002; Kawamura *et al*., 2006; Azimzadeh *et al*., 2008; Wright, Gallagher and Smith, 2009; Drevensek *et al*., 2012; Kirik, Ehrhardt and Kirik, 2012; Spinner *et al*., 2013; Kumari *et al*., 2021; Muroyama *et al*., 2023). Often, mutants with defects in PPB formation also have disrupted cortical microtubule organization, which may impede cell expansion (Torres-Ruiz and Jürgens, 1994; Whittington *et al*., 2001). An important study showed that the PPB is not critical for growth, as absence of >80% of PPBs generates macroscopically normal plants with relatively minor division plane orientation defects. Division plane positioning defects were attributed to spindle positioning defects while cortical microtubule organization appeared less affected (Ambrose and Cyr, 2008; Schaefer *et al*., 2017).

PPB formation requires the PROTEIN PHOSPHATASE TYPE 2A (PP2A) phosphatase B’’ regulatory subunit protein encoded by *fass/ton2* (Torres-Ruiz and Jürgens, 1994; Camilleri *et al*., 2002) in *A. thaliana* and two related genes in maize called *discordia1 (dcd1)* and *alternative discordia1 (add1)* (Gallagher and Smith, 1999; Wright, Gallagher and Smith, 2009). FASS/DCD1/ADD1 forms a complex with microtubule-binding proteins including TONNEAU1, TONNEAU1 RECRUITING MOTIF proteins, and other PP2A subunits that influence cortical microtubule organization and PPB formation (Wright, Gallagher and Smith, 2009; Spinner *et al*., 2013). *dcd1* mutants have ∼30% abnormal subsidiary cells, likely due to its high rate of defective PPBs (∼40%, examined by microtubule immunostaining) (Wright, Gallagher and Smith, 2009).

Here, we use the partially defective PPBs in *dcd1* single mutant to measure the contribution of PPB formation to division plane positioning. We also observe *de novo* recruitment of the division site protein TANGLED1 to misoriented cell plate fusion sites that is partially dependent on actin.

## Results and Discussion

### Defects in *dcd1* PPB formation result in reduction or lack of TAN1-YFP accumulation

To determine whether partially defective PPBs affect TAN1 recruitment to the division site, we observed the division site localized protein TANGLED1-YFP (TAN1–YFP) in *dcd1* and wild-type siblings with the microtubule marker CFP-TUBULIN (Martinez *et al*., 2017). Wild-type subsidiary cells had no defects in PPB formation or TAN1-YFP accumulation (n = 0/112 cells from 19 plants, Figure 1A, Fig. S1A). In contrast, *dcd1* mutant cells often had defective PPBs that incompletely encircled the cell, similar to previous results (Wright, Gallagher and Smith, 2009) (38%, n = 42/110 cells from 7 plants). Defective PPBs had uneven microtubule accumulation, including one-sided accumulation (“singular”, Figure 1B, Fig. S1B). Correspondingly, uneven or singular TAN1-YFP accumulation at the division site was observed in preprophase/prophase (35% n = 38/110 from 7 *dcd1* mutant plants) in metaphase and anaphase (35%, n = 16/46), and in telophase (41%, n =65/157, Figure 1C), suggesting that PPB establishment is required for TAN1 recruitment to the division site.

**Figure 1.**
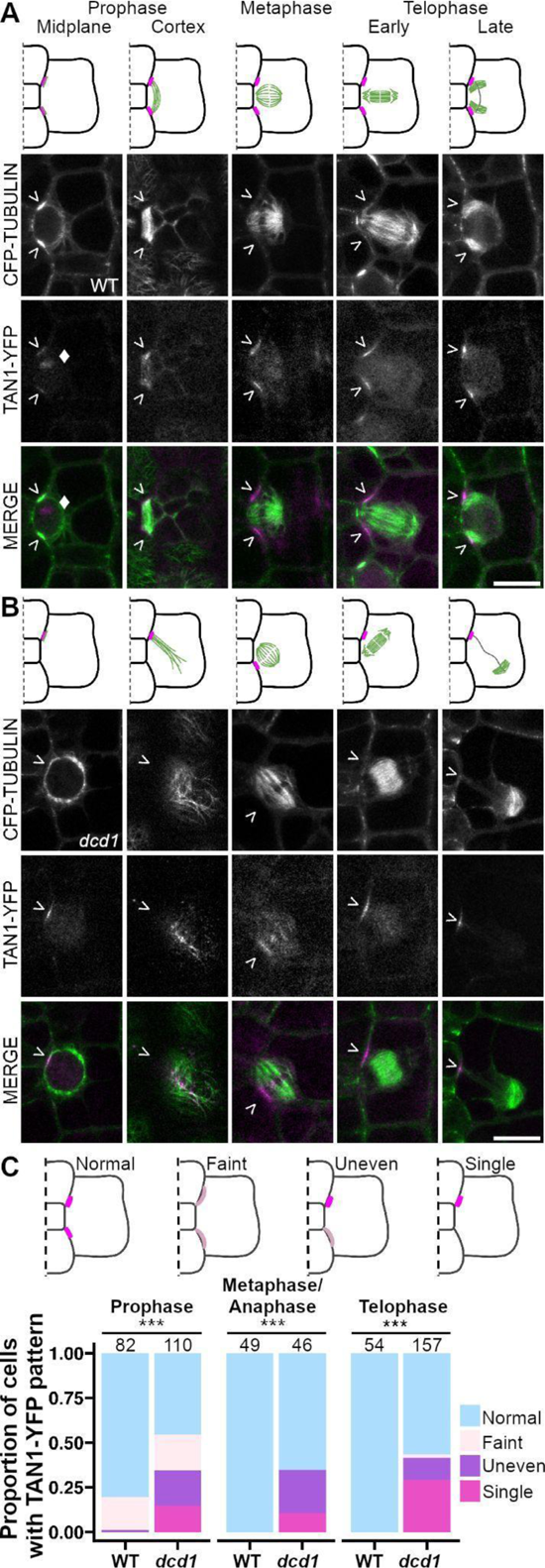
Both PPB formation and TAN1-YFP recruitment is defective in the *dcd1* mutant. (A-B) Model of (A) wild-type or (B) *dcd1* subsidiary cell divisions. Cell walls (black), microtubule structures (green), and TAN1-YFP (magenta) are shown. Below are representative images with CFP-TUBULIN labeling microtubules (green) and TAN1-YFP (magenta) labeling the division site (>) and sometimes the nucleolus indicated with a Diamond (♦). (C) Possible TAN1-YFP accumulation patterns. Darker and lighter shades of magenta represent higher and lower TAN1-YFP intensities reflecting more or less accumulation respectively. Below, stacked barplot comparing wild-type and *dcd1* cells that exhibit various TAN1-YFP patterns represented by the schematic models above. Numbers above bars represent cells examined. N = 19 wild-type plants and 7 *dcd1* plants. Scale bars = 10 μm.

### Defective PPBs in *dcd1* single mutants result in division plane positioning defects

Like many studies that have examined the role of the PPB in division plane positioning (Camilleri *et al*., 2002; Azimzadeh *et al*., 2008; Drevensek *et al*., 2012; Schaefer *et al*., 2017; W. Wang *et al*., 2019; Kumari *et al*., 2021), our initial analysis of *dcd1* mutants was performed using static images rather than time-lapse imaging. This data generated strong correlative support for the role of the PPB in division plane positioning, but actual cell division trajectories were not analyzed. To directly assess the relationship between PPB formation, TAN1 accumulation, and final division positioning, 12-minute time intervals were used to track divisions, capitalizing both on the invariant positions of subsidiary cell divisions and the *dcd1* mutant partial PPB formation defects (Figure 2A-D). At *dcd1* subsidiary cell division sites (n = 374 division sites total from 4 plants), we measured the TAN1-YFP and/or CFP-TUBULIN fluorescence intensities and classed final divisions as ‘oriented’ or ‘misoriented’ dependent upon whether the phragmoplast returned to the expected division site. We found that robust PPB microtubule accumulation strongly predicts correctly oriented cell divisions. Division sites with lower microtubule accumulation and undetectable TAN1-YFP tended to be misoriented (79%, n = 26/33 cells with TAN1-YFP fluorescence intensity at background levels, Figure 2E). For cell divisions captured in later stages, 94% (metaphase, anaphase, or telophase, n = 50/53) of misoriented final divisions were associated with undetectable TAN1-YFP intensity at the time lapse onset (Figure 2F, n = 112 cells). These data show that the PPB is essential for division plane positioning in subsidiary mother cell divisions and that TAN1-YFP localization at the division site is a reasonable proxy for previous preprophase band formation.

**Figure 2.**
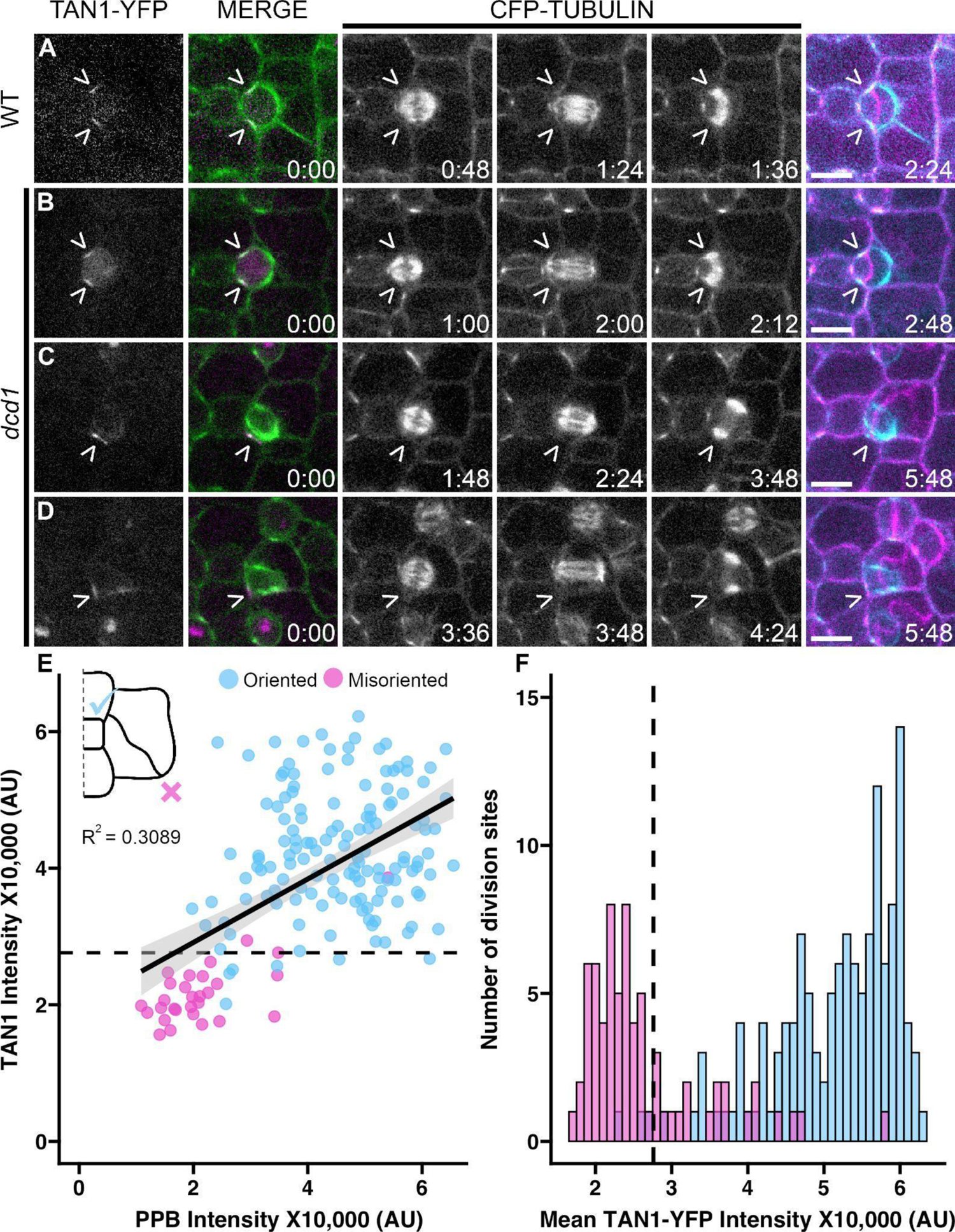
Defective preprophase bands and TAN1 localization result in misoriented divisions. Time lapses of subsidiary cell divisions expressing CFP-TUBULIN and TAN1-YFP in (A) wild-type cells and (B-D) *dcd1* cells. Left-most columns show TAN1-YFP localization at t = 0 (TAN1-YFP is magenta and microtubules are green in the merge). The last column overlays the the first (cyan) and last frames (magenta) of the microtubule channel showing the final division in relation to the preprophase band. Carets (>) mark the division site. Scale bars are 10 µm. (E) Comparative TAN1-YFP and PPB intensity from time lapses of *dcd1* cells. “Oriented” is when the phragmoplast returned to the expected division site and “misoriented” is when the cell plate inserted at atypical locations. n = 85 cells, N = 4 plants. (F) TAN1-YFP fluorescence intensity histograms colored by division orientation outcome (blue = oriented and magenta = misoriented) for time lapses that start after prophase in *dcd1*. The dotted line represents the visible detection limit. n = 112 cells. N = 4 plants.

### TAN1-YFP accumulates at misoriented cell plate insertion sites

*dcd1* phragmoplasts often direct cytokinesis to aberrant locations or *de novo* cell plate fusion sites. Interestingly, TAN1-YFP accumulated at *de novo* cell plate fusion sites (n = 21 misoriented phragmoplasts, N = 3 plants) (Figure 3A, B, E). Time-lapse imaging revealed that *de novo* TAN1-YFP accumulation trails behind the phragmoplast after it touches the cortex (Figure 3B, n = 22/22 cells from 3 plants, Fig. S1C). TAN1-YFP has been previously shown to accumulate near the phragmoplast midline (Martinez *et al*., 2017) This suggests that TAN1-YFP may be transported from the phragmoplast to the cell cortex, independently from the PPB.

**Figure 3.**
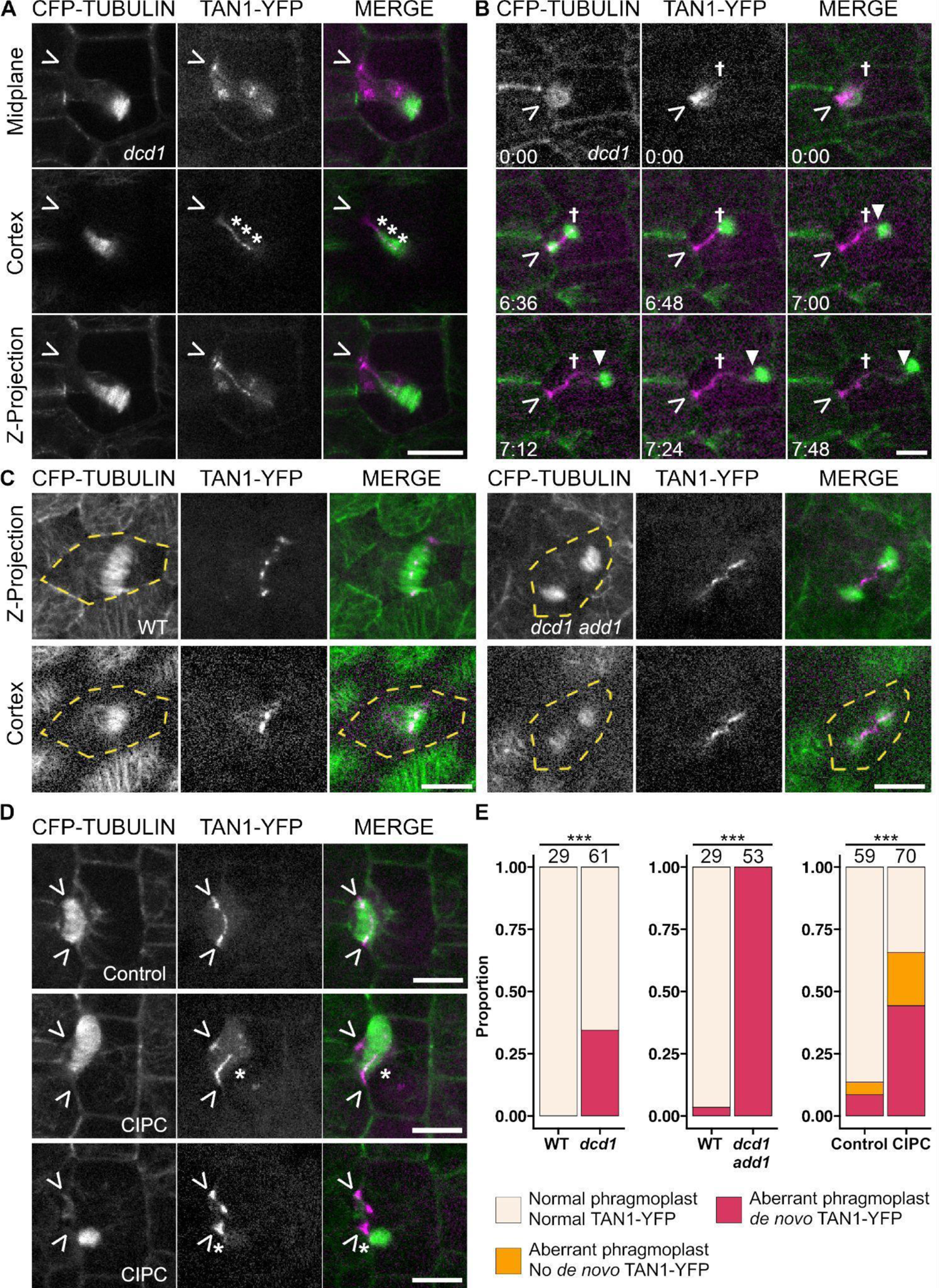
Cell plate insertion sites accumulate *de novo* TAN1-YFP. (A-D) CFP-TUBULIN (green) and TAN1-YFP (magenta) in various dividing cells. Carets (>) mark the division site and asterisks (*) mark *de novo* TAN1-YFP. (A) *dcd1* subsidiary mother cell with *de novo* cortex-localized TAN1-YFP indicated with asterisks. (B) Time lapse of a *dcd1* cell cortex during phragmoplast expansion. Dagger (✝) marks the edge of TAN1-YFP previously recruited in prophase and the triangle (▾) marks movement of the phragmoplast. Time stamps are in Hours:Minutes.(C) Z-projection and cortex views of wild type and *dcd1 add1* mutant embryos in telophase. Yellow dotted lines outline the cell. (D) Representative Z-projections of subsidiary mother cell phragmoplasts from CIPC and DMSO control treated samples. Asterisks mark *de novo* TAN1-YFP while carets mark the expected division site. (E) Barplots of *de novo* TAN1-YFP cell cortex accumulation in the *dcd1* mutant, *dcd1 add1* double mutant, or DMSO and CIPC treated wild-type plants. Numbers above bars represent total cell numbers. N ≥3 plants or kernels of each genotype or treatment. Asterisks indicate significant differences by Fisher’s Exact Test, P < 0.001. Scale bars = 10 µm.

TAN1-YFP was also observed at the cell cortex in the *dcd1 add1* double mutant cells that never make PPBs (Figure 3C, Fig. S2) (Wright, Gallagher and Smith, 2009). The *dcd1 add1* mutants are seedling lethal, so embryos were imaged 21 days after pollination. Wild-type cells showed normal TAN1-YFP accumulation at the division site at all stages (100%, n = 304 cells, n = 24 kernels, Fig. S2A). In the *fass/tonneau2* mutant and in cells treated with microtubule depolymerizing drugs, AtTAN::YFP was not observed at the cortex (Walker *et al*., 2007; Rasmussen, Sun and Smith, 2011). Similarly, in the *dcd1 add1* mutant, TAN1-YFP was not observed at the cortex in preprophase/prophase to anaphase cells (0%, n = 0/71 cells, n = 9 kernels, Fig. S2B). However, TAN1-YFP often accumulated at the cell cortex in telophase (72%, n = 36/50 cells from 9 kernels). Using higher resolution imaging, we found that TAN1-YFP accumulated only after the phragmoplast touched the cortex (100%, n = 53/53 cells, N = 4 kernels), but did not accumulate before (n = 11/11 cells from 4 kernels) (Figure 4C, Fig. S2). TAN1-YFP rarely accumulated at the cortex ahead of the phragmoplast (4%, n = 2/53, Fig. S3). These data further indicate that TAN1-YFP can be recruited to the cell cortex independently of the PPB.

**Figure 4.**
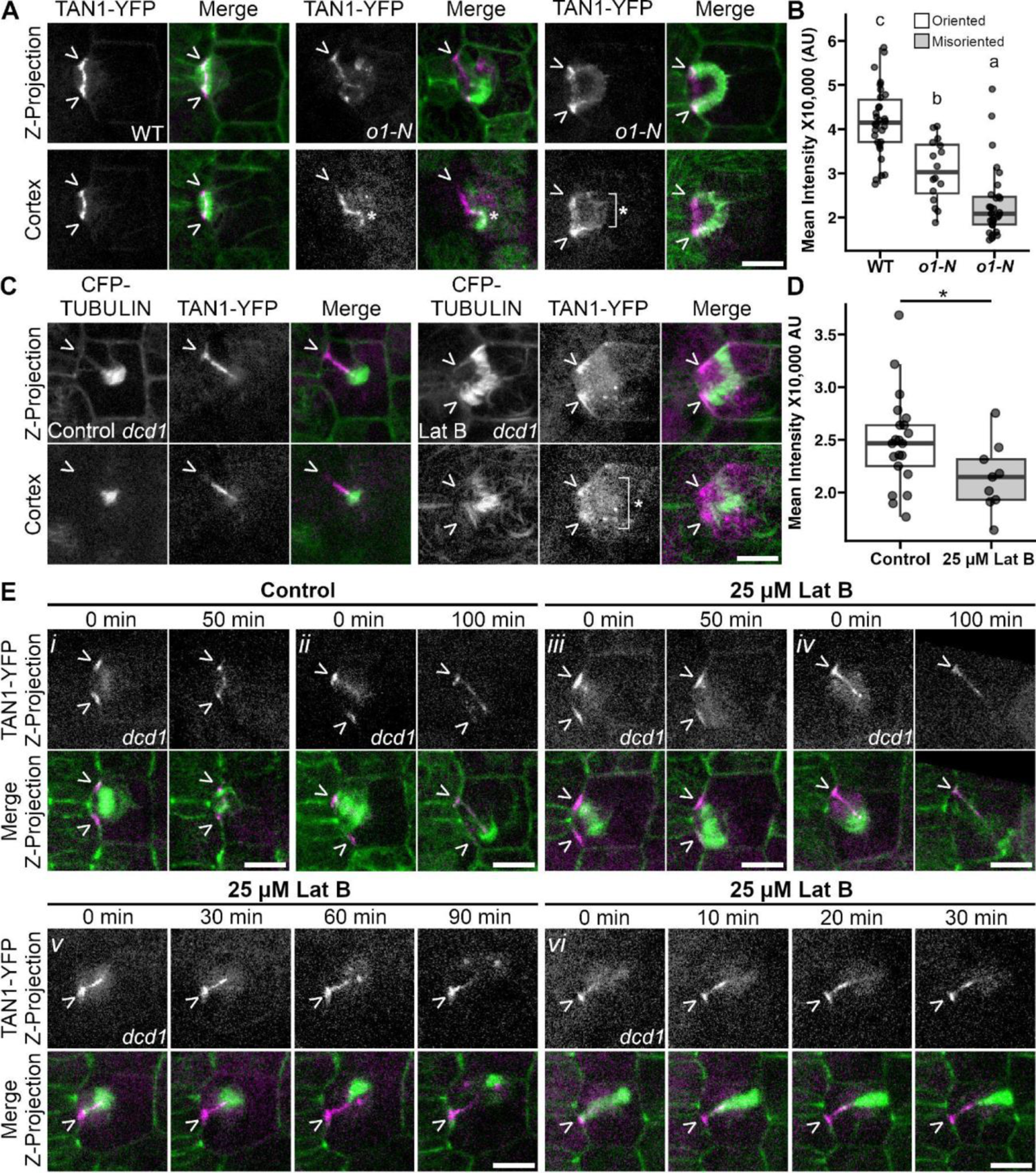
Actin and myosin XI motor protein OPAQUE1 increase TAN1 accumulation at *de novo* cell plate insertion sites. A) Subsidiary cell divisions in the *o1* mutant and wild-type siblings. B) Boxplot of TAN1-YFP intensities at telophase in oriented and misoriented divisions in wild type and *o1* mutant cells*. P* = 1.02e-12, One-way ANOVA followed by Tukey’s HSD, letters mark significant differences between groups. (C) TAN1-YFP accumulation in control and 25 µM Lat B treated *dcd1* cells. (D) Boxplot of TAN1-YFP intensity at misoriented divisions of *dcd1* in DMSO control (n = 23 cells, N = 2 plants) and 25 µM Lat B (n = 9 cells, N = 2 plants) treatments. P = 0.0417, Wilcoxon rank sum sum test. (E) Timelapse of *dcd1* cells in control and Lat B treatments. Carets (>) mark the division site and asterisks (*) mark *de novo* TAN1-YFP. Boxplot horizontal lines represent the quartiles and median. Whiskers are 1.5*IQR.

When additional or misoriented phragmoplast arms were generated in wild-type cells using the herbicide chlorpropham (CIPC), TAN1-YFP was also recruited to *de novo* cell plate fusion sites (Figure 3D). CIPC generates branched phragmoplasts through its tubulin binding activity but does not affect PPB formation (Liu, Joshi and Palevitz, 1995; Eleftheriou and Bekiari, 2000; Buschmann *et al*., 2006). Wild-type cells expressing *TAN1-YFP* and *CFP-TUBULIN* were treated for two hours with 0.7 µM or 1 µM CIPC or the respective DMSO controls and imaged. When additional or misoriented phragmoplast arms formed, *de novo* TAN1-YFP was observed after the phragmoplast contacted the cortex (Figure 3E, 67%, n = 31/46 cells from 3 plants).

### Actin and myosin XI OPAQUE1 facilitate TAN1-YFP accumulation at *de novo* cell plate insertion sites

We found that accumulation of TAN1-YFP at *de novo* cell plate insertion sites is partially dependent on the myosin XI O1. We hypothesized that since TAN1 interacts with PHRAGMOPLAST ORIENTING KINESIN1 (POK1) and POK2 (Müller, Han and Smith, 2006; Rasmussen, Sun and Smith, 2011; Mills, Morris and Rasmussen, 2022), and related kinesin 12s interact with myosin XI motor proteins including O1 (Huang *et al*., 2022; Nan, Liang, *et al*., 2023), O1 might be necessary for TAN1-YFP accumulation. TAN1-YFP fluorescence intensity during telophase was reduced but not absent in both correctly oriented and de novo cell plate fusion sites in *o1* compared to wild-type siblings (Figure 4A-B, p = 1.02e-12, One-way ANOVA followed by Tukey’s HSD). Therefore, O1 facilitates TAN1-YFP accumulation during telophase.

Actin filament disruption also reduced TAN1-YFP accumulation at *de novo* cell plate fusion sites. Since O1 likely moves along actin filaments, actin filament formation was inhibited with latrunculin B (Lat B) treatment in *dcd1* cells. 10-minute treatments with 25 µM Lat B inhibited actin polymerization (Fig. S4). Lat B treatment reduced TAN1-YFP accumulation at *de novo* cell plate fusion sites (Figure 4C-E, P = 0.0417, Wilcoxon rank sum sum test). To determine whether *de novo* TAN1-YFP recruitment or maintenance depends on actin filaments, 10-minute time points were taken after treating *dcd1* cells with control or 25 µM Lat B. In control-treated *dcd1* cells, TAN1-YFP accumulated and remained at the cell cortex as a narrow line following the phragmoplast trajectory (n = 15/17 cells, n = 4 plants). Rarely, TAN1-YFP accumulation was reduced (n = 1/17) or was not maintained at the cell cortex (n = 1/17). In Lat B treatments, TAN1 accumulation was often reduced (n = 13/18) or was not maintained after treatment (n = 5/18). Therefore, both TAN1-YFP recruitment and maintenance at *de novo* sites are reduced when actin filaments were disrupted.

In the absence of PPB-mediated recruitment, we observe TAN1-YFP accumulation at *de novo* cell plate fusion sites that is partially dependent on actin filaments and O1. Consistently, when actin is disrupted in Arabidopsis root cells, TAN1, POK1 and myosin XI division-site localization becomes diffuse (Huang *et al*., 2022). Actin has been shown to connect the leading edge of the phragmoplast with the division site through the action of myosin VIII in *Physcomitrium patens* (Wu and Bezanilla, 2014) and is required for division plane positioning (Mineyuki and Palevitz, 1990; Gallagher and Smith, 1999; Frank, Cartwright and Smith, 2003; Gilliland *et al*., 2003; Galatis and Apostolakos, 2004; Facette and Smith, 2012; Vaškebová, Šamaj and Ovecka, 2017). During the late stages of phragmoplast expansion, actin is needed for the completion of cell plate fusion (van Oostende-Triplet *et al*., 2017), potentially in part dependent on recruitment of TAN1 and other division site localized proteins. Recruitment of other division site proteins (e.g POK2) to *de novo* cell plate fusion sites have also been observed in mutants which generate additional ectopic cell plates, suggesting that *de novo* localization may be a common feature during cytokinesis (Lebecq *et al*., 2023).

TAN1-YFP is recruited to the cell cortex independent of the PPB, by the phragmoplast. Further, actin filaments and the myosin O1 both contribute to TAN1-YFP recruitment during late telophase. We hypothesize that TAN1-YFP accumulation may reflect the assembly of entire “division-site modules”, which may have yet unknown roles in late cytokinesis.

## Acknowledgments

We thank Jennifer Phan and other Rasmussen Laboratory members for help with the generation and propagation of genetic resources. We thank Dr. Meng Chen for the Zeiss 880 (NIH 3R01GM087388-06S2). For Zeiss 880 training, we thank Dr. Marschal Bellinger (UCR) and Dr. David Carter (UCR Microscopy and Imaging Core Facilities).

## Author Contributions

**ANU**: Conceptualization, Investigation, Formal Analysis, Visualization, Writing - Original Draft, Writing - Review & Editing, Project administration, Funding acquisition; **BND**: Methodology, Investigation, Formal Analysis, Visualization, Funding acquisition; **LAA**: Methodology, Project administration; **SGG**: Methodology, Investigation, Formal Analysis; **SM**: Investigation, Formal Analysis, Funding acquisition; **JJK**: Formal Analysis; **SA**: Formal Analysis, Funding acquisition; **CGR**: Conceptualization, Resources, Supervision, Funding acquisition, Writing - Original Draft, Writing - Review & Editing, Project administration, Funding acquisition.

## Data availability

All relevant data can be found in the article and its supplementary information.

## Materials and Methods

### RESOURCE AVAILABILITY

#### Lead contact

Further information and requests for resources and reagents should be directed to and will be fulfilled by Carolyn Rasmussen.

#### Materials availability

This study did not generate new unique reagents.

#### Data and code availability

- All data reported in this paper will be shared by the lead contact upon request.
- This paper does not report original code.
- Any additional information required to reanalyze the data reported in this paper is available from the lead contact upon request.

#### EXPERIMENTAL MODEL DETAILS

Maize (*Zea mays*) plants were grown in standard greenhouse conditions (31-33°C temperature setpoints with supplementary lighting from 5-9 PM at ∼400 µ E m^-2^ s^-1^) in 1 L pots with soil (20% peat, 50% bark, 10% perlite, and 20% medium vermiculite) supplemented with additional magnesium nitrate (50 ppm N and 45 ppm Mg) and calcium nitrate (75 ppm N and 90 ppm CA) and Osmocote Classic 3-4M (NPK 14-14-14 %, AICL SKU#E90550). Alternatively, plants were grown in the field (Agricultural Operations https://agops.ucr.edu/, Riverside, CA, USA) to generate maize embryos, which were hand harvested from ears 21-23 days after pollination.

## METHOD DETAILS

### Plant material and genotyping/phenotyping

Plants expressing CFP-ß-TUBULIN and/or TAN1-YFP (Mohanty *et al*., 2009; Wu *et al*., 2013) were genotyped with CFP-TUBULIN forward primer GFP5FOR (5’-GCGACGTAAACGGCCACAAGTTCAG-3’) and the reverse primer TubB3433R (5’-CGGAAGCAGATGTCGTAGAGC-3’) and the TAN1-YFP forward primer TAN LSP1 (5’-ACGACCGTTAGCACAGAACC-3’) and the reverse primer GFP5Rev (5’-CTGAACTTGTGGCCGTTTACGTCGC-3’), or identified by painting leaves with 4 g/L glufosinate (Finale, Bayer) in 0.1% Tween 20 (Sigma). Resistance to glufosinate was assessed after 2–5 d.

The *dcd1 add1* and *dcd1* mutants were a kind gift from Dr. Amanda Wright. The *dcd1-mu1* and *add1* alleles were genotyped according to Wright et al. 2009 (Wright, Gallagher and Smith, 2009) using the forward MuE2 (5’-TCCATAATGGCAATTATCTC-3’) and the reverse 55862nrev (5’-GGTGCTACATATACGCTAAAG-3’) for dcd1-mu1 and the forward 3dCAPbfor (5’-GTTGTTTTCCCCCTTGGATT-3’) and the reverse 3dCAPbrev (5’-CTTGAGTTCTTGTTTGCTCAG-3’) for *add1*. To distinguish between wild type and *add1* mutant alleles, PCR products were digested with the restriction enzyme KpnI overnight and then run on a 4% agarose gel for 90 minutes at 110V. *dcd1* mutant plants were also identified by phenotype using glue impressions of epidermal leaf cells (Allsman, Dieffenbacher and Rasmussen, 2019). The *opaque1/dcd2* mutants were a kind gift from Dr. Michelle Facette. *o1-N1242A* mutants were identified by phenotype using a lightbox and/or glue impressions.

Leaves were dissected for imaging after 3-5 weeks of growth from the whorl until the ligule was 2 mm from the base and the abaxial epidermal cells were placed into a Rose chamber as described (Rasmussen, 2016) to observe dividing cells. For live imaging of wild-type and *dcd1 add1* double mutant embryos, maize plants were grown in the greenhouse or in the field under standard conditions. Ears were harvested 21-23 days after pollination. Embryos were dissected from kernels and loaded onto a Rose chamber with the flat plumule face down (Kiesselbach, 1949).

### Chemical treatments

1 M CIPC (CAS 101-21-3 from TCI, #C2555) was dissolved in DMSO. Leaf samples were loaded in 0.7 µM or 1 µM CIPC or the respective 0.07% or 0.1% DMSO control in a rose chamber and imaged after 1 to 2 hours of treatment. Samples were loaded into 25 µM Lat B (Fisher Scientific, #2182-1) or the respective DMSO control. Z stacks were acquired 2 hours after treatment. For time lapse imaging, samples were loaded directly into 40 µl of 25 µM Lat B and a time lapse was started with 10 minute time points. To identify what concentration of Lat B was required to depolymerize actin filaments, leaf tissue samples were treated with 0.0025 µM, 0.25 µM, or 25 µM Lat B for 1 hour, fixed, and stained with Alexa Fluor 488 Phalloidin (Fisher Scientific, #A12379) following Nan *et al*. 2019 (Nan, Mendoza and Facette, 2019).

### Confocal Microscopy

Micrographs and time-lapse data were acquired using a Yokogawa W1 spinning disk microscope with an EM-CCD camera (Hamamatsu 9100c) on a Nikon Eclipse TE inverted stand. Solid-state Obis lasers with power ranging from 40 to 100 mW were used in combination with standard emission filters (Chroma Technology). For TANGLED1-YFP, a 514 nm laser with emission filter 540/30 nm was used. For CFP-TUBULIN, a 445 nm laser with emission filter 480/40 nm was used. Oil or water immersion objectives (60X/1.2 NA, 100X/1.45 NA) were used. Images and time-lapses were taken with Micromanager-1.4 using a 3-axis DC servo motor controller and ASI Piezo Z stage. For time-lapse, 10 or 12 minute time intervals were used as specified with Z-intervals ranging from 3 to 5 µm. For Z-stacks acquired with no timelapse, 0.5 µm steps were used.

Images were also acquired using a Zeiss LSM 880 confocal laser scanning microscope (100X oil objective immersion lens, NA = 1.46) with Airyscan super resolution mode and Z-intervals of 0.25 µm or 3 µm. The 0.25 µm Z-intervals were used to generate the X-Z projection in Fig. S1C. A 514 nm-excitation laser with bandpass filters 465-505 with long-pass 525 filter was used. Images were processed using default Airyscan settings with Zen software (Zeiss).

### Figure Preparation

Figures were made using Gnu Image Manipulation Program (Gimp, version 2.10.32, https://www.gimp.org/). Image levels were only adjusted linearly and images were enlarged or rotated with no interpolation.

### Accessions

CFP-TUBULIN and TAN1-YFP lines were generated by the Maize Cell Genomics Group (Mohanty *et al*., 2009). Gene sequences can be found at MaizeGDB (https://www.maizegdb.org/gbrowse) using the following accession numbers (B73, v4): *DISCORDIA 1* (Zm00001d024857), *ALTERNATIVE DISCORDIA 1* (Zm00001d010862), and *TANGLED 1* (Zm00001d038060).

## QUANTIFICATION AND STATISTICAL ANALYSIS

Time lapse images, X-Z projections, and Z-projections were generated using Fiji (ImageJ, http://rsb.info.nih.gov/ij/, RRID:SCR_003070). Mean fluorescence intensity was measured using the “straight” or “oval” tool. X-Y drift in time lapses was corrected using the translation function in the StackReg plug-in in ImageJ (Thévenaz, 1998) or the Fast4DReg plugin (Laine *et al*., 2019). Analysis of TAN1-YFP localization and/or intensity measurements was done by separating the CFP-TUBULIN channel from the TAN1-YFP channel and using the CFP-TUBULIN channel to identify the stage of cell division and location at the midplane or the cell cortex.

For Figure 1, TAN1-YFP localization to the division site was described as “Normal”, “Faint”, “Uneven”, or “Single” based on the presence or absence of localization and TAN1-YFP intensity at the cell midplane. “Normal” intensity describes wild type TAN1-YFP localization– two bright accumulations in the subsidiary mother cell that flank the guard mother cell. “Faint” describes two accumulations that are less intense than “normal”. “Uneven” describes two accumulations, one that is more intense than the other. Finally, “Single” describes cells with TAN1-YFP accumulation at one division site and absence from the other.

For the TAN1-YFP mean intensity measurements in Figure 2E and F, a line ROI was drawn at the cell midplane, bisecting the region of TAN1-YFP accumulation. For cells in prophase, the same ROI was used to measure CFP-TUBULIN accumulation in the preprophase band at the division site (Figure 2E). When TAN1-YFP or CFP-TUBULIN accumulation was below detection as frequently observed in *dcd1* subsidiary mother cell divisions, the ROI was selected at the expected division site location for a subsidiary mother cell division.

When analyzing *de novo* TAN1-YFP localization in *dcd1*, *dcd1 add1*, or the CIPC treated cells in Figure 3E, phragmoplasts were categorized as normal or aberrant, where aberrant includes misoriented phragmoplasts and split phragmoplasts in the CIPC treatments (Figure 3E). TAN1-YFP localization was determined to be “normal” if TAN1-YFP was only observed to localize to the division site, and “*de novo*” if TAN1-YFP was observed to accumulate at *de novo* cell plate fusion sites, which were identified by observing the phragmoplast and the cell cortex.

For cortical TAN1-YFP intensity measurements in Figure 4B and D, mean intensity was measured using a 2 µm line ROI. For misoriented phragmoplasts, ROIs were drawn starting from the leading edge of the phragmoplast along the phragmoplast midline.

Graphs, tables, and statistics were generated using R(R Core Team, 2023) and Rstudio (Posit team, 2023) using the following packages: tidyr, ggplot2, ggprism, ggpubr (Wickham, 2016; Dawson, 2022; Kassambara, 2023; Wickham *et al*., 2023; Wickham, Vaughan and Girlich, 2023). Statistical details of experiments can be found in the main text and/or figure legends.

Significance was defined as P < 0.05 and parametric tests were used unless data distribution was non-normal, whereupon an equivalent non-parametric test was used instead. In Figure 4B, the One-way ANOVA was followed by a Tukey’s HSD multiple comparison test. For the comparison of categorical variables in Figure 1C and Figure 3E, a Fisher’s exact test was used.

## KEY RESOURCES TABLE

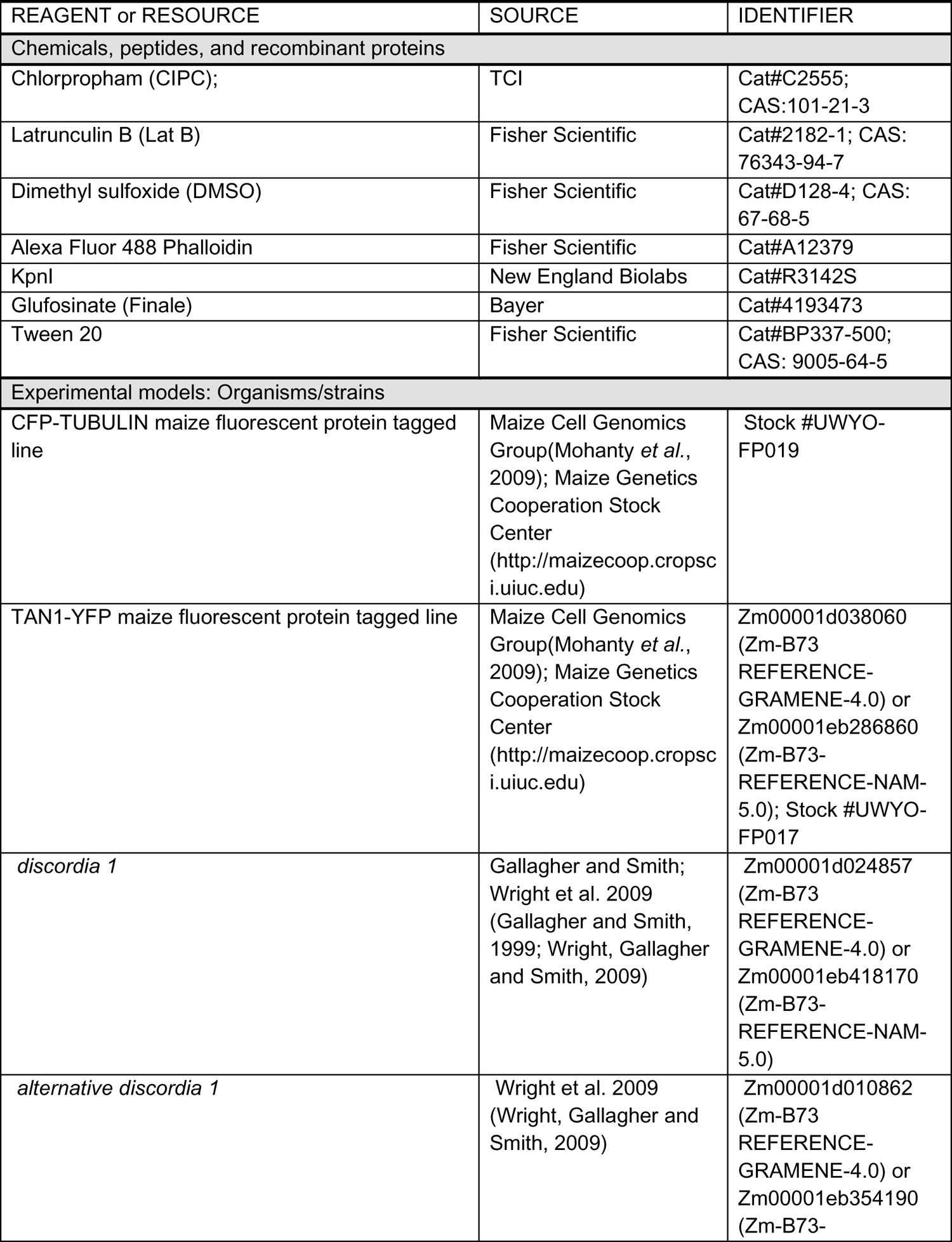
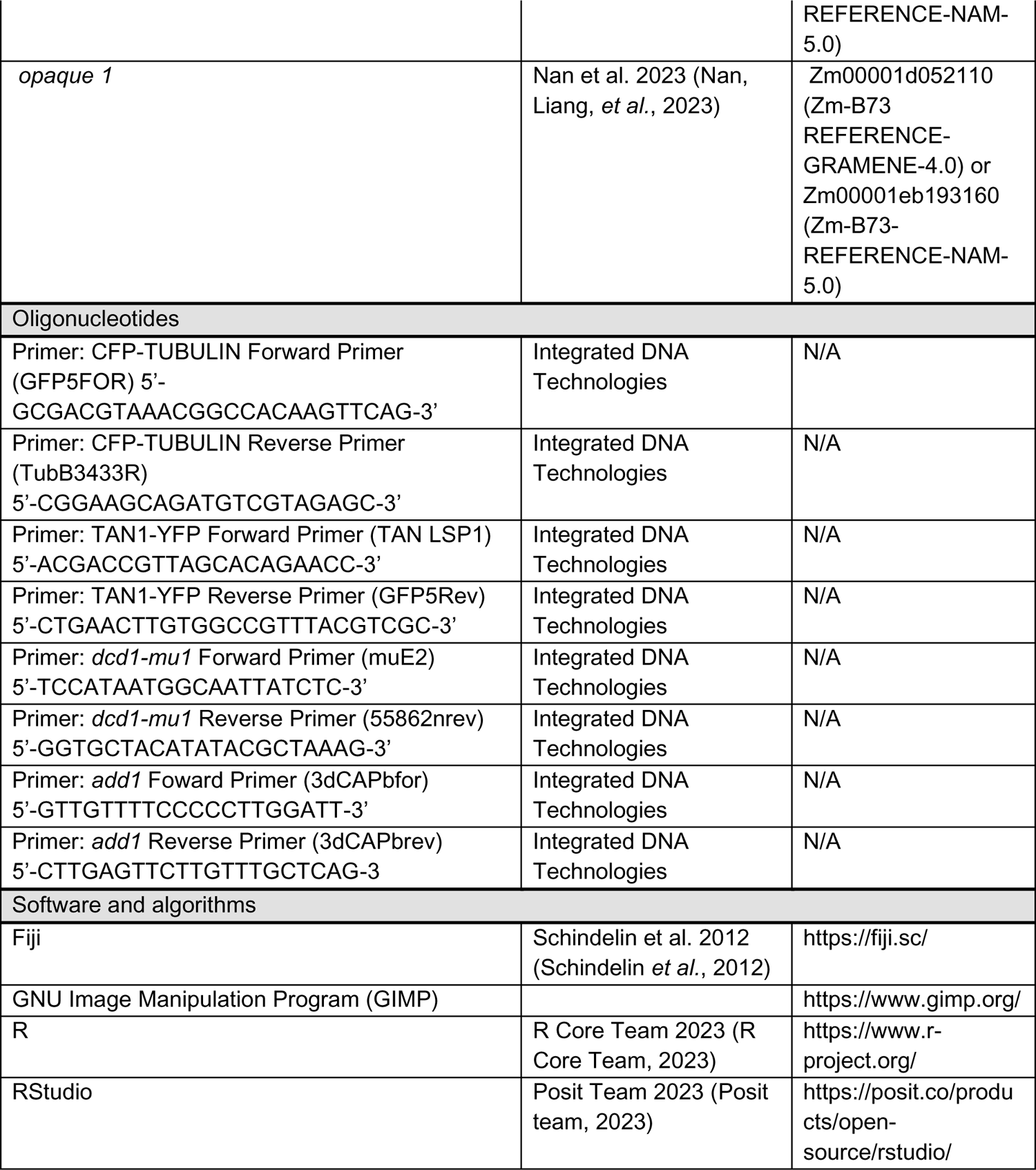

---

**Fig. S1.**
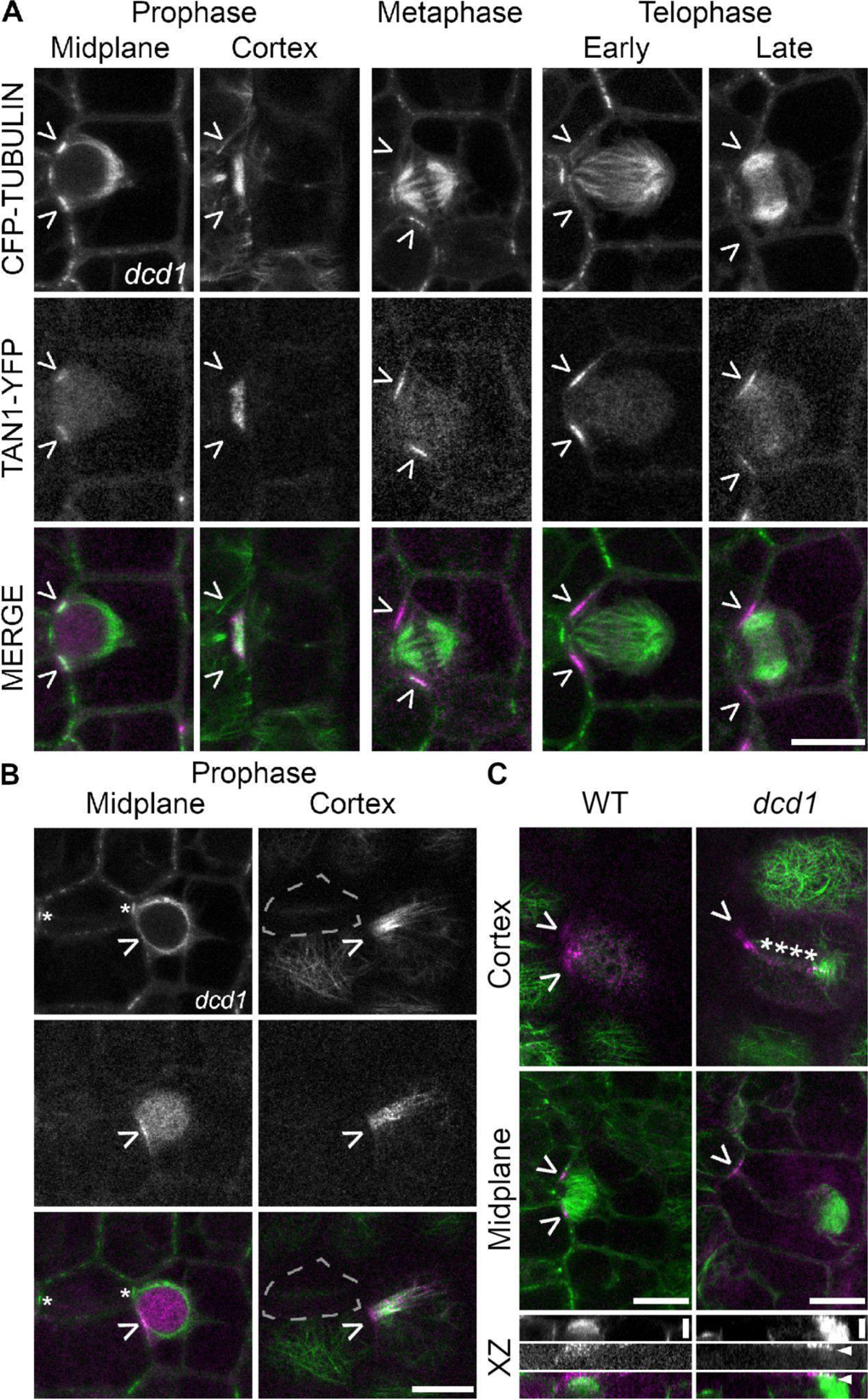
Confocal micrographs of divisions in wild type (WT) and *dcd1* plants. (A) Representative images of correctly oriented *dcd1* divisions expressing CFP-TUBULIN (green) and TAN1-YFP (magenta) with typical microtubule structures and TAN1-YFP localization. (B) An additional example of a defective preprophase band in *dcd1* with CFP-TUBULIN and TAN1-YFP accumulation on one division site and missing from the other. Asterisks mark the typical interphase microtubule accumulation in the neighboring guard mother cell. Dotted lines outline the guard mother cell. Carets point to the division site. (C) Micrographs of wild type (left) and *dcd1* (right) cells in telophase at the cell cortex and midplane expressing CFP-TUBULIN (microtubules, green) and TAN1-YFP (magenta). Below, the CFP-TUBULIN, TAN1-YFP, and merged channels of XZ-projections showing the side view of the cell. Z-slices were taken at 0.25 µm intervals. Scale bars for A-C cortex and midplane view are 10 µm, and 3.4 µm for the XZ projections in C.

**Fig. S2.**
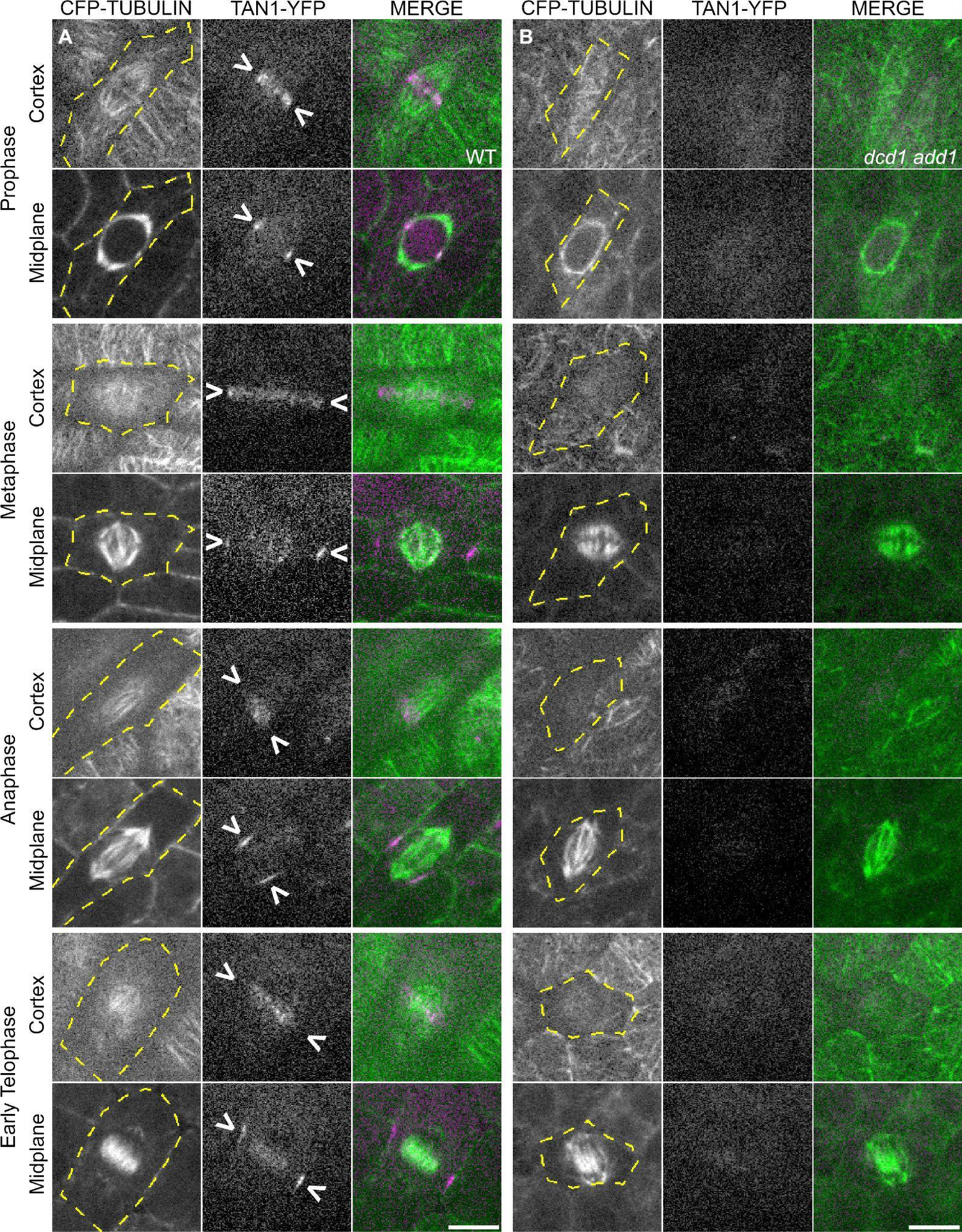
TAN1-YFP localization in wild type and the *dcd1 add1* double mutant from prophase to early telophase. Micrographs of cortex and midplane views of (A) wild-type embryos and (B) *dcd1 add1* embryos expressing CFP-TUBULIN (microtubules, green) and TAN1-YFP (magenta). Arrowheads point to TAN1-YFP localization to the division site and a yellow dotted line marks the cell outline. Scale bars are 10 µm, all images in the two panels are the same magnification.

**Fig. S3.**
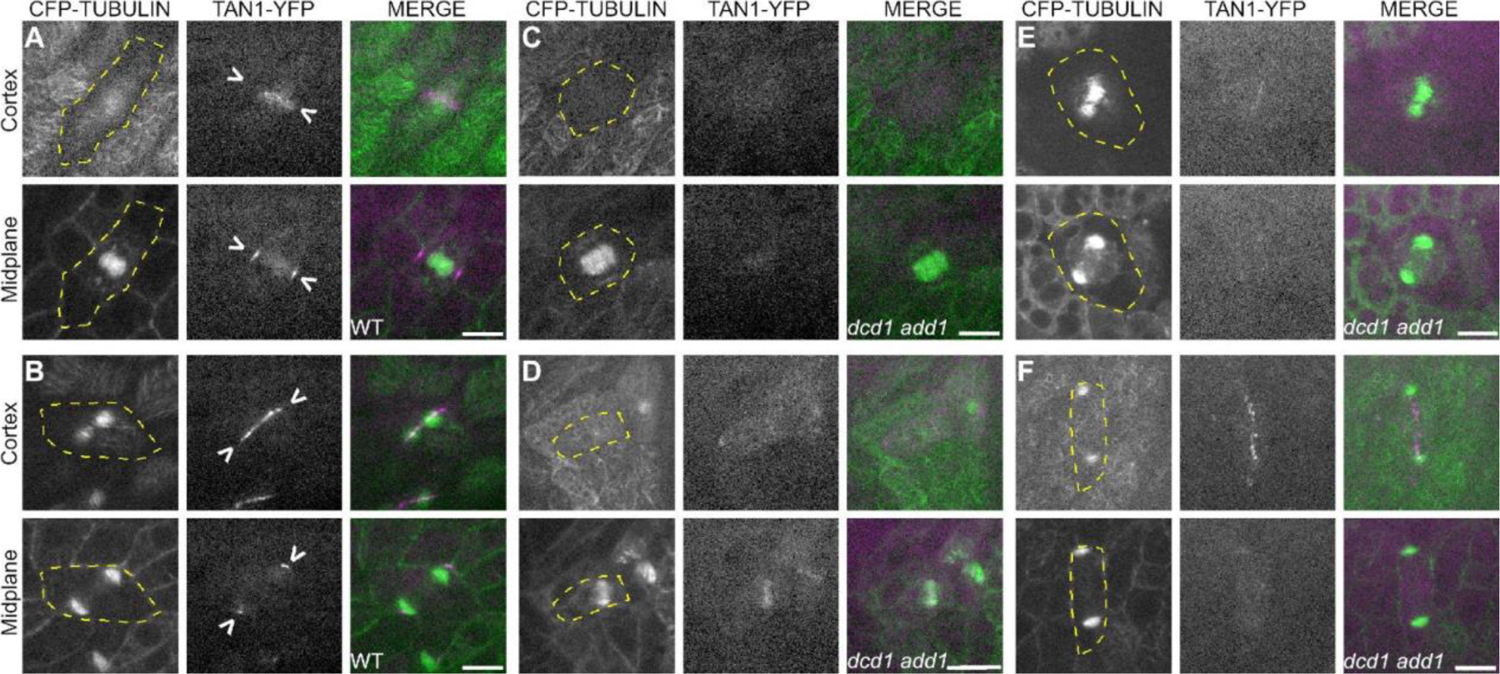
TAN1-YFP localization patterns in *dcd1 add1* embryos. (A-B) TAN1-YFP localizes to the division site (>) in wild-type embryos (A) before and (B) after the phragmoplast has reached the cell cortex. (C-D) In early telophase cells in *dcd1 add1*, TAN1-YFP is (C) absent from or (D) diffuse at the cell cortex before the phragmoplast has fully expanded. TAN1-YFP is also visible in the phragmoplast midline. (E-F) In late telophase cells in *dcd1 add1*, (E) TAN1-YFP localizes to the cell cortex as a narrow band once the phragmoplast reaches the cortex and (F) rarely localizes ahead of phragmoplast expansion. Scale bar is 10 µm.

**Fig. S4.**
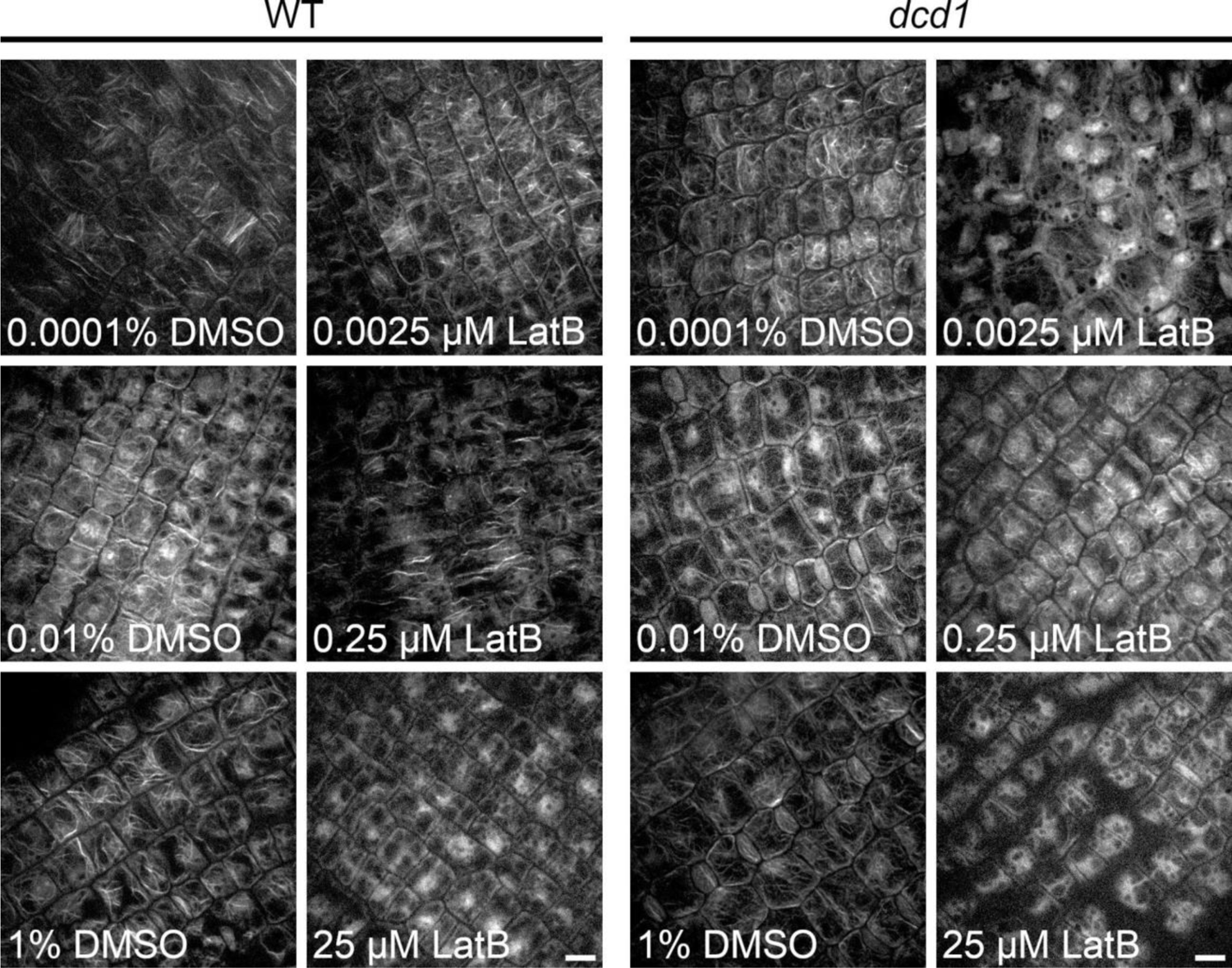
Optimization of latrunculin B treatment for *dcd1* and its wild-type sibling. Micrographs of actin filaments immunostained with Alex fluor 488-phalloidin. Scale bar is 10 µm.

